# Evidence that Nek1 does not phosphorylate Rad54-S572 during recovery from IR

**DOI:** 10.1101/2022.04.01.486731

**Authors:** Ishita Ghosh, Md Imtiaz Khalil, Arrigo De Benedetti

## Abstract

A main focus of the work in our lab is on the activity of Tousled Like Kinase 1 (TLK1) in the area of DNA Damage and Repair. As one of its key interactor, TLK1 phosphorylates NIMA related kinase 1 (Nek1), and Nek1 was reported by Spies et al. to phosphorylate and regulate the activity of the key HRR protein Rad54^1^, which suggested an intriguing signal transduction pathway: TLK1>Nek1>Rad54. In an effort to confirm such relations, we now report that we have not been able to reproduce key findings from that study. Specifically, we found that Nek1 does not phosphorylate RAD54-S572 as was reported. We generated Nek1-KO mouse NT1 cells^2^ and Nek1-Knock-down in Hek293 (same cells as in Spies et al.), and the pRAD54-S572 signal does not change with our custom Ab, with or w/o IR. When we used an Ab from the Lobrich lab, it detected an immunoreactive band of wrong size for RAD54, which also did not change after IR even in synchronized G2 cells, contrary to their report. We also note that their P-assignment was based on guessing a weak consensus Nek1 sequence, and that site-directed mutagenesis of RAD54-S572 failed to yield biological effects in their in in vitro studies^1^. To conclusively establish that S572 is not a site of phosphorylation of Nek1, we carried out a IVK with purified Nek1 and RAD54 followed by MS analysis of the phosphatides, which revealed several but not S572. We also could not reproduce their copurification of Nek1-RAD54 by coIP, calling into question this interaction. Neither we could reproduce their results demonstrating the importance of Nek1 for HRR using the same SceI-mediated DR-GFP conversion assays.

## INTRODUCTION

Each day our cells face many exogenous and endogenous challenges that threat the genomic stability. The DNA double strand break is the most lethal form of damage. The major mechanism to maintain the fidelity of genetic code is Homologous recombination repair (HRR). The central enzyme of this pathway is Rad54.^3^ Rad54 is a well conserved protein of the Rad52 group, a generally well conserved group in eukaryotes. Rad54 is a motor protein that translocates along dsDNA in an ATP hydrolysis dependent manner.^4^ In mammals, the RAD54 gene plays its most important role during early developmental stages.^5^ The primary function of homologous recombination in mitotic cells is to repair double-strand breaks (DSBs) that form as a result of replication fork collapse, from processing of spontaneous damage, and from exposure to DNA-damaging agents like topoisomerase poisons.^6^ Ideally known as the swiss-army knife of homologous recombination repair, Rad54 functions in different stages of HRR. Rad54 interacts with Rad51 and promotes DNA strand -exchange, translocates along donor template, remodels donor chromatin, promotes Branch-migration of Holliday Junctions.^7–10^ With such multi-faceted role in vivo, any post-translational modification of Rad54 may alter its activity. Therefore, it is critical to understand the regulation of this error-free mode of DNA repair in cells.

## RESULTS

Given the possibility of a TLK1>Nek1>Rad54 axis and its importance for HRR, we wanted to probe the pattern of pRad54-S572 in cells with perturbed TLK1 activity. In an initial communication with Marcus Lobrich, he stated that they had finished the original antiserum published in the Mol Cell paper but that they would ship us a replacement that was not well-characterized. We tested that antiserum during a time-course of recovery from IR of HEK293 cells, either control or expressing a dominant Nek1 kinase-hypoactive mutant (HEK N5)^11^, which was expected to show reduced pRad54-S572 signal (Fig.1); and it was immediately obvious that there were problems. The main immunoreactive band ran on the gel faster than its expected position of ~80kDa for Rad54 (Fig1). The signal was not very clean as there were several other (presumably less specific) prominent bands. Most importantly, the intensity of the pRad54 did not increase after recovery from IR, as it was reported. Given the impossibility of reproducing that part of the work with the Ab they provided, we commissioned our own custom-made pRad54-S572 antiserum from Thermofisher. After the 3d boost, an ELISA showed that the serum had a high titer (>2X10^5^ dilution) and an affinity >100-fold for the phosphopeptide compared to unphosphorylated, even before affinity purification. To establish the specificity of the Ab and its dependence on Nek1, we carried out a siRNA-mediated knockdown of Nek1 in Hek293 cells, similarly to their Fig. 3D experiment. Despite the successful knock-down of Nek1 (Fig.2), we did not observe any change in the signal with our pRad54-S572 antiserum, which detected a band at the correct position for Rad54 and overlapping the signal obtained with a commercial (santa-cruz sc-166370) pan-Rad54 antiserum. This clearly indicated that Nek1 is unlikely to be the kinase responsible for phosphorylation of S572. Considering the possibility that there might still be sufficient Nek1 in the KD-cells to phosphorylate S572, we took advantage of a mouse PCa cell line in which we clonally knocked-out Nek1 via CRISPR/CAS9 to rule out such possibility. Surprisingly, there was no immunoreactive band with pRad54-S572 antiserum in either control or the Nek1-KO clone – note that these cells do express Rad54L and Rad54B (RNAseq data communicated by Xiuping Yu). In contrast, there was a strong band in HEK293, equally in control and Nek1-siRNA-KD cells (Fig. 3). This result suggests that despite the high sequence conservation between mouse and human (Fig. 3B), the pRad54-S572 is not found in mouse, and one should wonder whether such proposed phosphorylation is at all important. To probe this question further, we set out to establish if their report of Rad54-S572 phosphorylation after IR, and particularly in late G2, could be reproduced. We thus, synchronized Hek293 cells following release from G1/S block with HU and irradiated them in early G2 (Fig.4), repeating their experimental conditions. Contrary to what they reported, we found no increase in pRad54-S572 in either asynchronous or cells enriched G2 (Fig.5), for which 8h was reported in their paper as the maximal level of phosphorylation^1^.

**Figure 1.**
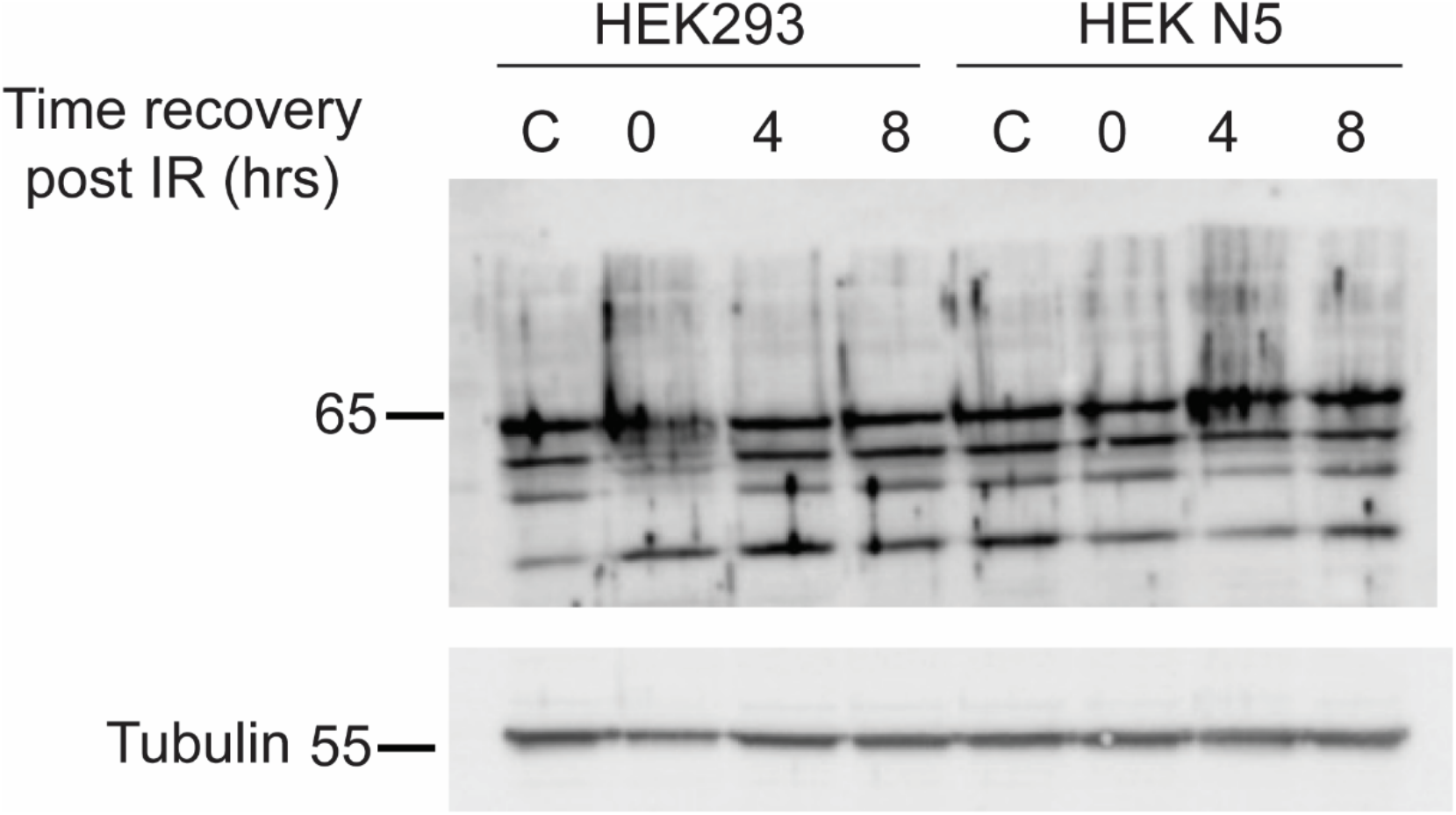
2X 10^6 HEK293 cells were seeded 24hrs prior to treatment and then cells were treated with 10Gy of I.R and allowed to recover for indicated times (hrs). 40μg of untreated control (C) and samples were loaded in 8% SDS-PAGE gel and probed with aliquot of antibody sample from Lobrich lab. Tubulin probed as loading control.

**Figure 2.**
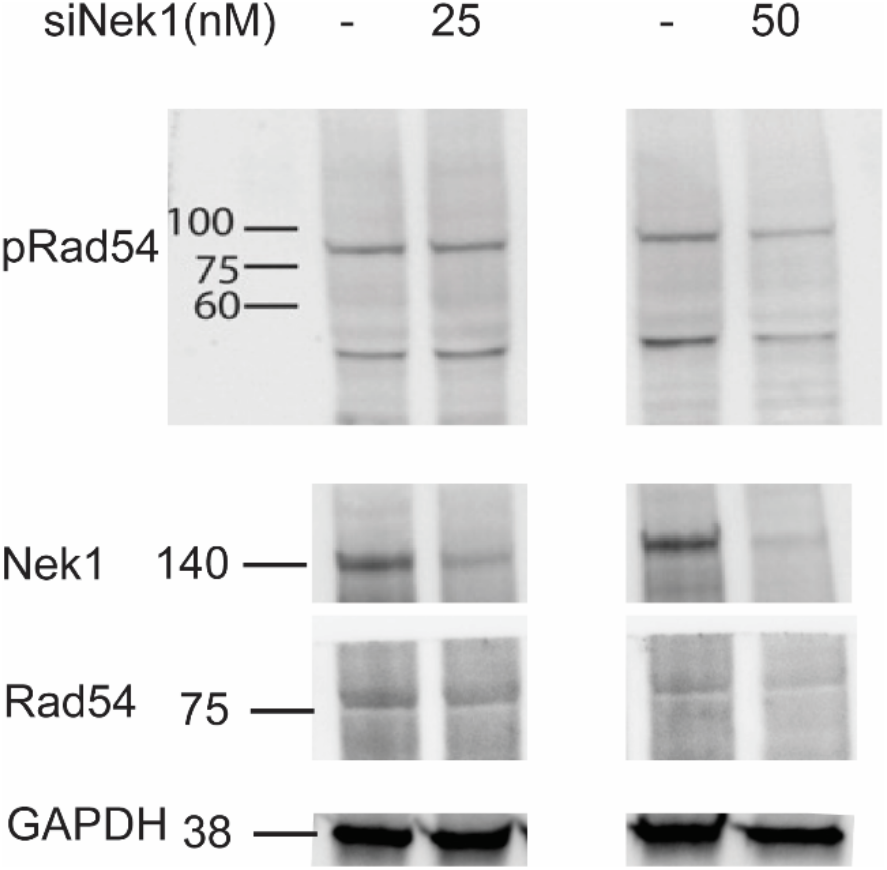
0.3X10^6 HEK293 cells seeded in 6 well plate 24hrs prior to siRNA treatment. Lipofectamine 3000 was used as transfection reagent with indicated amount of siNek1. 40ug of total samples loaded in 8% SDS-PAGE gel and probed with indicated antibodies (Nek1-sc 398813; Rad54-sc 166370; pRad54-Custom generated-Thermofisher; GAPDH-CST #2118)

**Figure 3.**
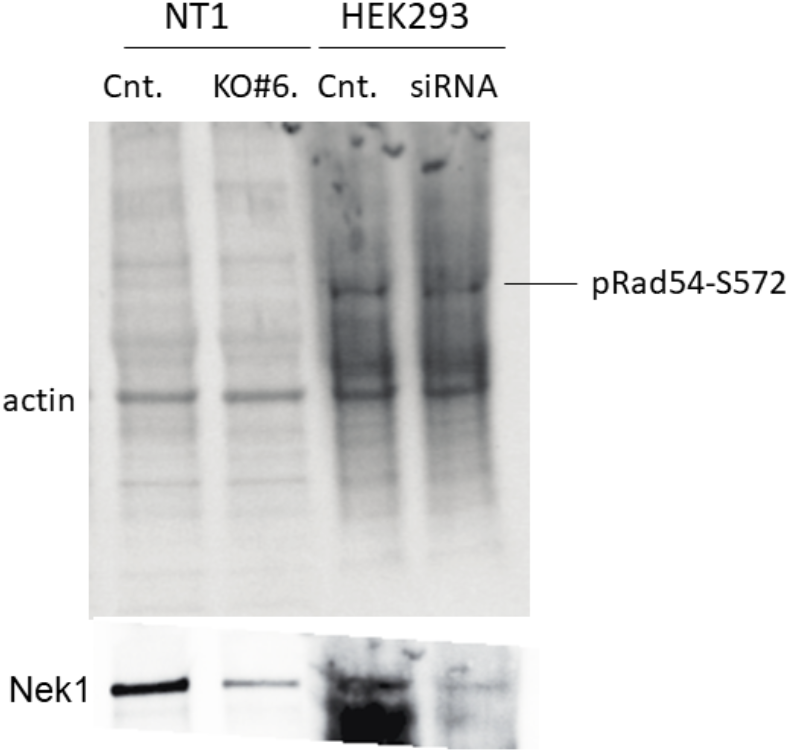
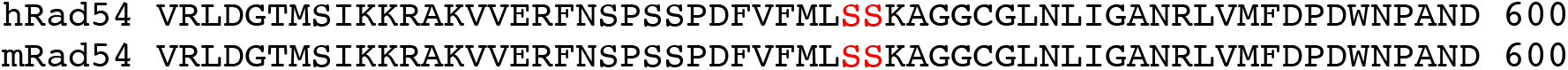
A) 40μg of cell lysate from NEK1 WT (Cnt), K.O #6 clone (in NT1 cells) and siRNA treated HEK293 cells and control (Cnt) loaded in 8% SDS-PAGE gel and probed for Nek1 and pRad54 (custom generated) antibody. Actin band shown as loading control. B) Human Rad54 and mouse Rad54 protein sequence alignment shown for the conservation of the S572 site.

**Figure 4.**
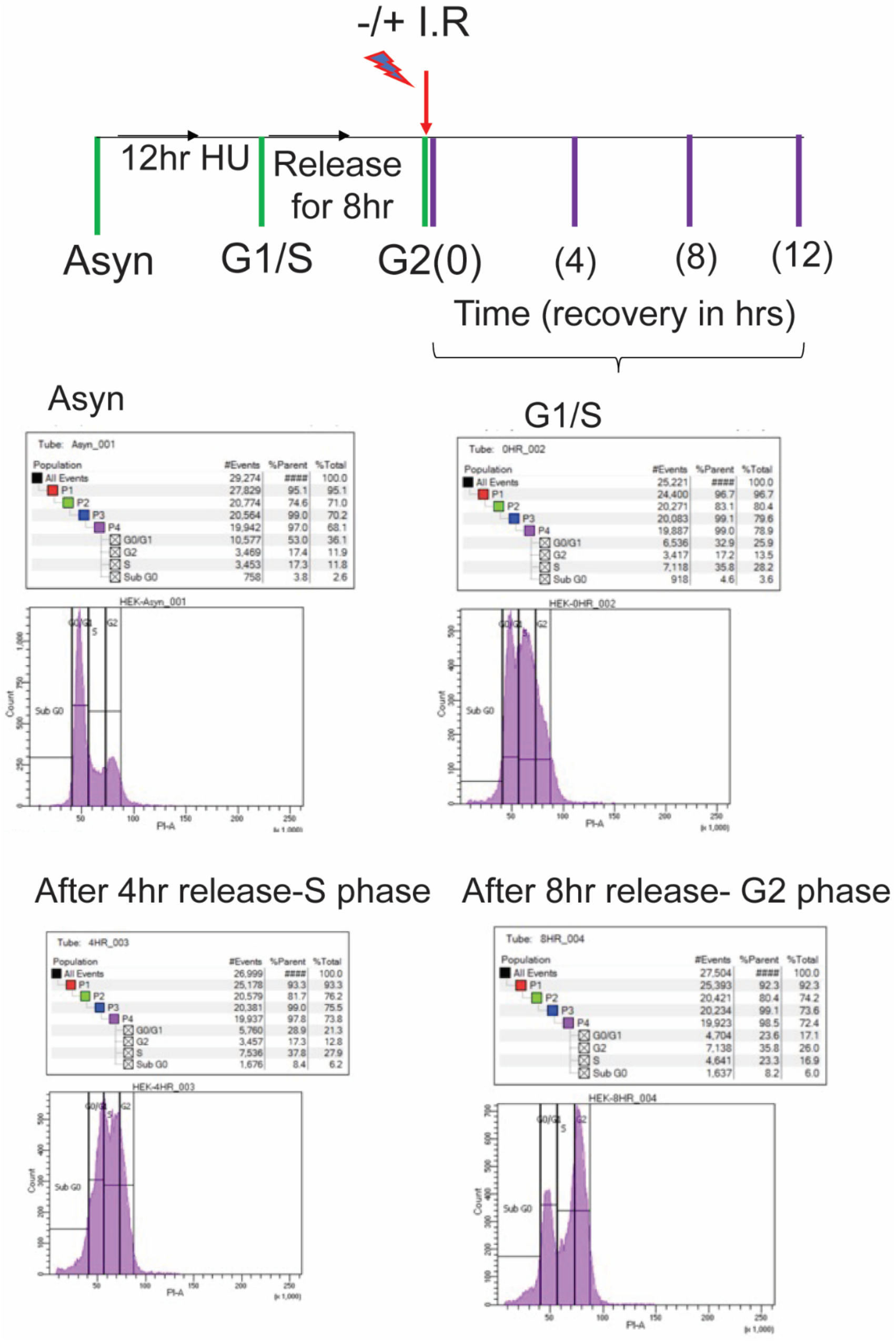
Cell cycle analysis of HEK293 cells synchronized (asynchronous -Asyn) at G1/S with Hydroxyurea treatment and released for indicated times (as shown in schematic).

**Figure 5.**
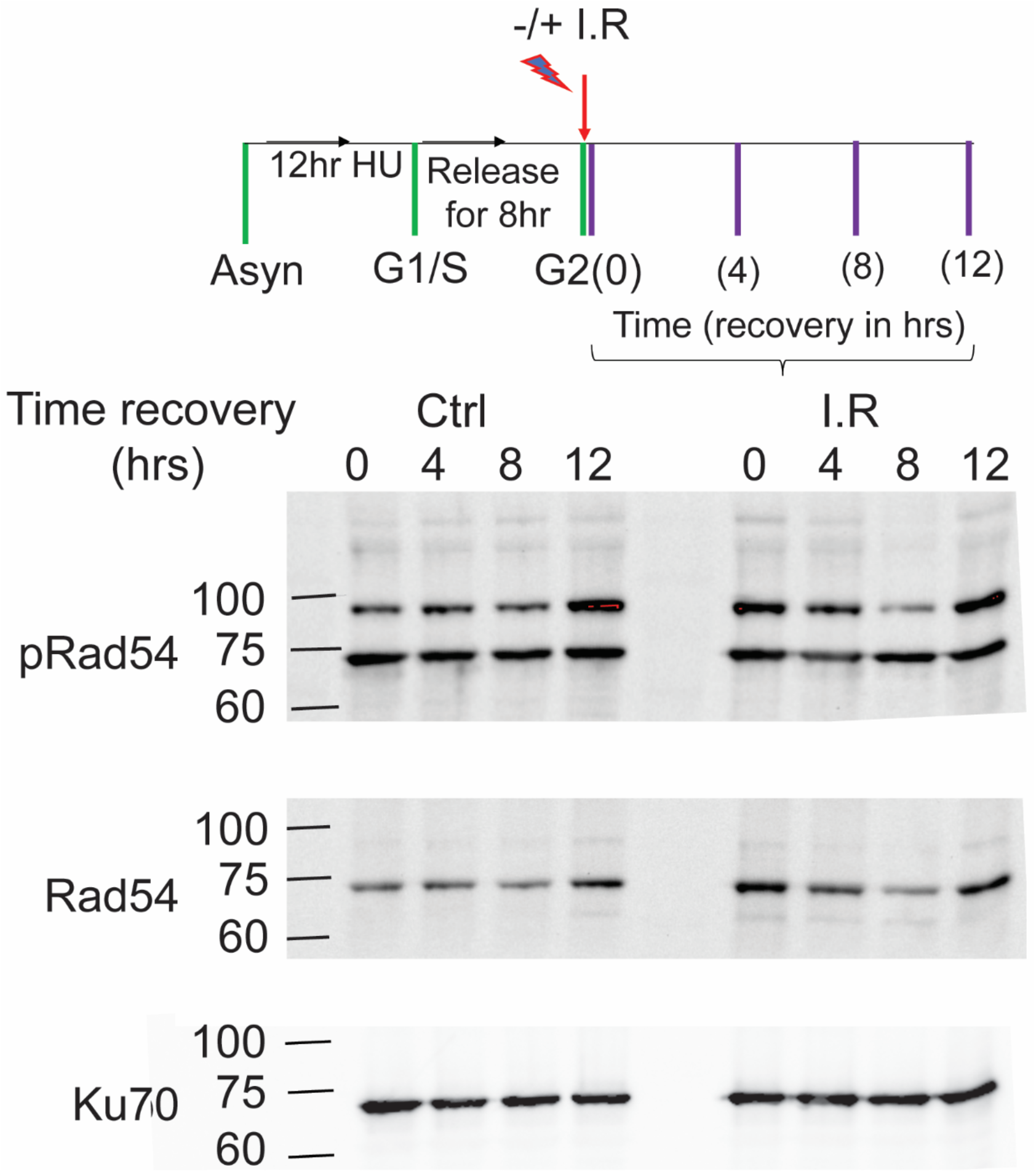
Synchronised and aynchronised HEK293 cells were untreated or irradiated (5 Gy) at G2 and allowed to recover for indicated times. 40ug of cell lysate was loaded in 8% SDS-PAGE gel and probed with indicated antibodies. Ku-70 probed for loading control.

In order find truly unbiased evidence for the proposed phosphorylation of S572 by Nek1, we carried out an *in vitro* kinase (IVK) and MS phosphopeptides analysis of recombinant (and functional) RAD54 phosphorylated by highly purified Nek1, as described in^12^. Note that a IVK reaction does not imply that RAD54 in a true in vivo substrate of Nek1, but this was done (Fig. 6A) merely to replicate the conditions used by Spies et al. (Fig 3B) in order to assign the putative phosphorylated residue. Note that the only possible interpretation of their Fig 3B is that S572 is the primary, if not the only, phosphor-target of Nek1, since virtually all P32 labeling from the IVK reaction is lost with the GFP-RAD54-S572/A mutant protein. In complete disagreement with their published Fig.3B and their assumed phosphorylation of S572 as the only Nek1 target site, there were several residues phosphorylated in vitro – but not S572 (Fig. 6B). A complete list of all modified the peptides is shown (Sup Table 1), and multiple Ser, Thr, and Tyr were found – note that Nek1 is a dual specificity kinase^13^. Note also that our preparation of Nek1 was fully active, as indicated also by its autophosphorylation at Y315 (MS data not shown), which is diagnostic of its level of activity. In conclusion, their assignment of S572 as the only phosphorylation site by Nek1 cannot be reproduced when analyzed correctly by MS instead of relying on a putative consensus sequence; and their results of Fig.3B remains very difficult to explain.

**Figure 6.**
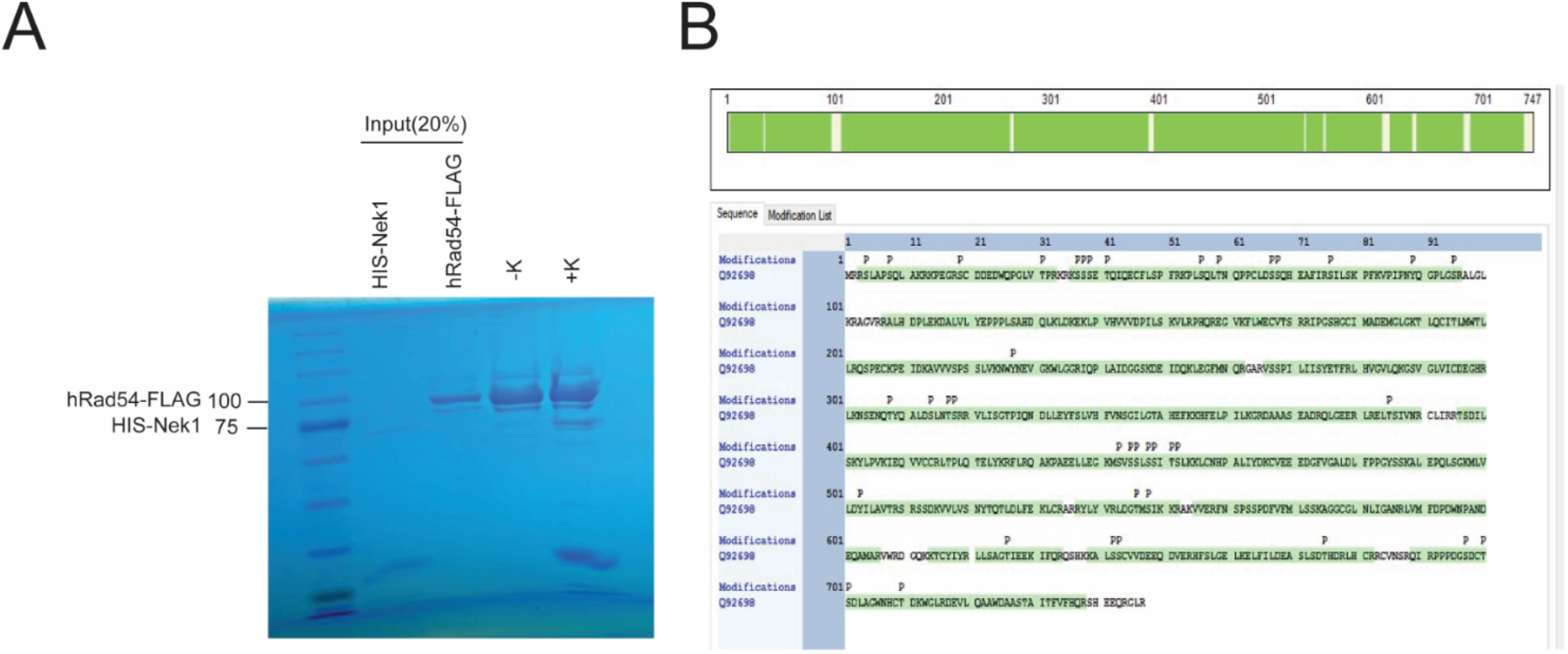
Determination of the RAD54 phosphopetides obtained by IVK with purified Nek1. A) Picture of the Coomassie stained gel confirming the purity of the proteins is shown. B) Phospho-peptide map of purified human Rad54 protein in IVK assay with purified Nek1.

Finally, we were not able to reproduce the reported interaction between Nek1 and Rad54 by coIP, although we cannot rule out that this could be due to a technical failure. Nonetheless, key findings of their paper could not be reproduced. These include a failure to attribute the phosphorylation of S572 to Nek1, its dependency on IR-induced damage, and in general the significance of such Rad54 modification for HRR.

To establish if Nek1 has any significant role in HRR, we carried out SceI-mediated DR-GFP conversion assays in Hek293 cells overexpressing siRNA-resistant wt-Nek1 or the Nek1-T141A dominant negative mutant (HEK N5), similar to the work reported in Fig. 4 of the Spies et al. paper. Again, contrary to their findings, we found that wt-Nek1 did not stimulate HRR and that the N5 overexpressing cells showed insignificant effects on HRR (Fig. 7). Depletion of endogenous Nek1 with siRNA did not change the picture, demonstrating that it is not the endogenous Nek1 that misrepresents the fact that neither wt nor a dominant hypomorphic mutant of Nek1 has any effect on HRR, in contradiction of the results reported in Fig.4 of the Spies et al paper for the S572A overexpressing mutant. In summary, we were not able to reproduce any of the key results of their paper about the importance of Nek1 for HRR.

**Figure 7.**
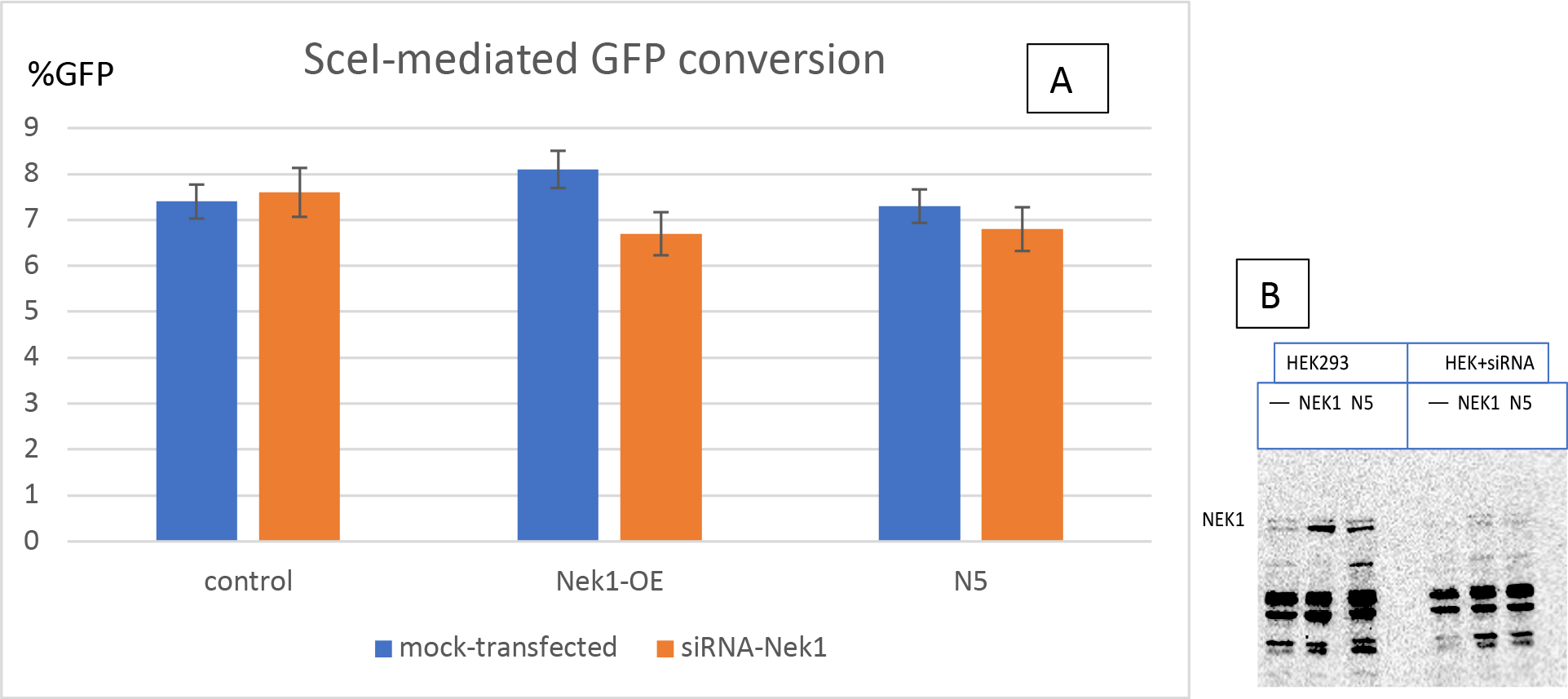
A. SceI-mediated DR-GFP conversion assays. Hek293 were stably transfected with a DR-GFP cassette and then with wt-Nek1 or dominant-negative Nek1-T141A (N5) expression constructs. HRR assays were then carried out 48h after transfection of SceI by FACS analysis of the proportion of GFP converted cells. B. Expression of wt-Nek1 or dominant-negative Nek1-T141A. Where indicated, siRNA to endogenous Nek1 was included 24h earlier (demonstration of the efficacy of the knock-down was already published in^11^).

## Acknowledgements

We thank Xinngui Shen (ORCID Xinggui Shen http://orcid.org/0000-0002-4002-967) of the Mass Spectrometry Core at LSU Health-Shreveport supported by LSU Health Shreveport Foundation, Center for Cardiovascular Diseases and Sciences, and Feist-Weiller Cancer Center. This work was supported by a grant from the Feist Weiller Cancer Center from LSUHSC, Shreveport, and DoD grant PC160398.

---

**Supplemental Table 1.**

| Confidence | Sequence | Modification | Quality PEP | Quality q-value | # Protein G | # Proteins | # PSMs | Master Protein |
| --- | --- | --- | --- | --- | --- | --- | --- | --- |
| High | LVMFDPDV | 2xOxidatio | 3.95E-05 | 0.004739 | 1 | 1 | 79 | Q92698 |
| High | IEQVVCCR | 2xCarbami | 0.000465 | 0.004739 | 1 | 1 | 40 | Q92698 |
| High | TLQCITLMV | 2xCarbami | 0.103627 | 0.004739 | 1 | 1 | 1 | Q92698 |
| High | KLCNHPALI | 2xCarbami | 2.21E-07 | 0.004739 | 1 | 1 | 59 | Q92698 |
| High | LCNHPALIY | 2xCarbami | 4.37E-07 | 0.004739 | 1 | 1 | 97 | Q92698 |
| High | QIRPPPDG | 2xCarbami | 4.81E-06 | 0.004739 | 1 | 1 | 5 | Q92698 |
| High | QIRPPPDG | 2xCarbami | 9.26E-11 | 0.004739 | 1 | 1 | 38 | Q92698 |
| High | YLPVKIEQV | 2xCarbami | 0.0002 | 0.004739 | 1 | 1 | 58 | Q92698 |
| High | MLVLDYILA | 1xPhospho | 0.186682 | 0.004739 | 1 | 1 | 1 | Q92698 |
| High | NWYNEVGK | 1xPhospho | 0.025781 | 0.004739 | 1 | 1 | 4 | Q92698 |
| High | LLSAGTIEEK | 1xPhospho | 0.0006 | 0.004739 | 1 | 1 | 1 | Q92698 |
| High | LRELTSIVNF | 1xPhospho | 0.128289 | 0.004739 | 1 | 1 | 1 | Q92698 |
| High | ELFILDEASL | 1xPhospho | 0.005986 | 0.004739 | 1 | 1 | 2 | Q92698 |
| High | SLAPSQLAK | 1xPhospho | 0.0191 | 0.004739 | 1 | 1 | 6 | Q92698 |
| High | RSLAPSQLA | 1xPhospho | 0.00741 | 0.004739 | 1 | 1 | 2 | Q92698 |
| High | SILSKPFK | 1xPhospho | 0.001382 | 0.004739 | 1 | 1 | 3 | Q92698 |
| High | VPIPNYQGF | 1xPhospho | 0.025547 | 0.004739 | 1 | 1 | 4 | Q92698 |
| High | SILSKPFKVF | 1xPhospho | 0.000224 | 0.004739 | 1 | 1 | 16 | Q92698 |
| High | LDGTMSIK | 1xPhospho | 0.010477 | 0.004739 | 1 | 1 | 3 | Q92698 |
| High | LKNSENQTY | 1xPhospho | 0.003824 | 0.004739 | 1 | 1 | 9 | Q92698 |
| High | LKNSENQTY | 1xPhospho | 0.011858 | 0.004739 | 1 | 1 | 8 | Q92698 |
| High | MSVSSLSSI | 1xPhospho | 0.003098 | 0.004739 | 1 | 1 | 12 | Q92698 |
| High | NSENQTYQ | 1xPhospho | 0.000117 | 0.004739 | 1 | 1 | 5 | Q92698 |
| High | NSENQTYQ | 1xPhospho | 0.000167 | 0.004739 | 1 | 1 | 16 | Q92698 |
| High | LVMFDPDV | 1xOxidatio | 2.01E-05 | 0.004739 | 1 | 1 | 90 | Q92698 |
| High | LDGTMSIK | 1xOxidatio | 0.001572 | 0.004739 | 1 | 1 | 39 | Q92698 |
| High | LDGTMSIK | 1xOxidatio | 0.00119 | 0.004739 | 1 | 1 | 15 | Q92698 |
| High | LEGFMNQR | 1xOxidatio | 0.003824 | 0.004739 | 1 | 1 | 27 | Q92698 |
| High | IQPLAIDGG | 1xOxidatio | 1.81E-06 | 0.004739 | 1 | 1 | 82 | Q92698 |
| High | MLVLDYILA | 1xOxidatio | 4.03E-06 | 0.004739 | 1 | 1 | 108 | Q92698 |
| High | MSVSSLSSI | 1xOxidatio | 4.98E-05 | 0.004739 | 1 | 1 | 22 | Q92698 |
| High | MSVSSLSSI | 1xOxidatio | 2.69E-06 | 0.004739 | 1 | 1 | 57 | Q92698 |
| High | FLRQAKPAE | 1xOxidatio | 0.090318 | 0.004739 | 1 | 1 | 13 | Q92698 |
| High | FNSPSSPDF | 1xOxidatio | 0.000102 | 0.004739 | 1 | 1 | 52 | Q92698 |
| High | DEIDQKLEG | 1xOxidatio | 0.003576 | 0.004739 | 1 | 1 | 8 | Q92698 |
| High | ALEPQLSGK | 1xOxidatio | 0.035362 | 0.004739 | 1 | 1 | 1 | Q92698 |
| High | EGVKFLWE | 1xCarbami | 0.000689 | 0.004739 | 1 | 1 | 3 | Q92698 |
| High | RIPGSHGCI | 1xCarbami | 7.72E-07 | 0.004739 | 1 | 1 | 48 | Q92698 |
| High | RIPGSHGCI | 1xCarbami | 2.86E-07 | 0.004739 | 1 | 1 | 69 | Q92698 |
| High | GSVGLVICD | 1xCarbami | 0.000188 | 0.004739 | 1 | 1 | 198 | Q92698 |
| High | GSVGLVICD | 1xCarbami | 0.00034 | 0.004739 | 1 | 1 | 1 | Q92698 |
| High | RIPGSHGCI | 1xCarbami | 6.88E-08 | 0.004739 | 1 | 1 | 110 | Q92698 |
| High | IPGSHGCIN | 1xCarbami | 0.000937 | 0.004739 | 1 | 1 | 22 | Q92698 |
| High | IPGSHGCIN | 1xCarbami | 6.54E-06 | 0.004739 | 1 | 1 | 37 | Q92698 |
| High | IPGSHGCIN | 1xCarbami | 1.5E-08 | 0.004739 | 1 | 1 | 19 | Q92698 |
| High | KALSSCVVD | 1xCarbami | 1.58E-05 | 0.004739 | 1 | 1 | 14 | Q92698 |
| High | ALSSCVVDE | 1xCarbami | 0.002291 | 0.004739 | 1 | 1 | 2 | Q92698 |
| High | ALSSCVVDE | 1xCarbami | 7.16E-06 | 0.004739 | 1 | 1 | 25 | Q92698 |
| High | ALSSCVVDE | 1xCarbami | 0.000106 | 0.004739 | 1 | 1 | 3 | Q92698 |
| High | FLWECVTSF | 1xCarbami | 6.98E-05 | 0.004739 | 1 | 1 | 162 | Q92698 |
| High | FLWECVTSF | 1xCarbami | 0.033493 | 0.004739 | 1 | 1 | 1 | Q92698 |
| High | QSPECKPEI | 1xCarbami | 9.62E-05 | 0.004739 | 1 | 1 | 36 | Q92698 |
| High | QSPECKPEI | 1xCarbami | 6.08E-05 | 0.004739 | 1 | 1 | 17 | Q92698 |
| High | TLQCITLMV | 1xCarbami | 8.25E-05 | 0.004739 | 1 | 1 | 119 | Q92698 |
| High | AGGCGLNL | 1xCarbami | 2.13E-05 | 0.004739 | 1 | 1 | 22 | Q92698 |
| High | AGGCGLNL | 1xCarbami | 0.008985 | 0.004739 | 1 | 1 | 1 | Q92698 |
| High | TLQCITLMV | 1xCarbami | 3.74E-06 | 0.004739 | 1 | 1 | 152 | Q92698 |
| High | KLCNHPALI | 1xCarbami | 8.56E-06 | 0.004739 | 1 | 1 | 68 | Q92698 |
| High | KTCYIYR | 1xCarbami | 0.000716 | 0.004739 | 1 | 1 | 9 | Q92698 |
| High | SCDDEDWC | 1xCarbami | 0.01714 | 0.004739 | 1 | 1 | 6 | Q92698 |
| High | LCNHPALIY | 1xCarbami | 4.64E-05 | 0.004739 | 1 | 1 | 131 | Q92698 |
| High | SCDDEDWC | 1xCarbami | 2.62E-06 | 0.004739 | 1 | 1 | 20 | Q92698 |
| High | TCYIYR | 1xCarbami | 0.002952 | 0.004739 | 1 | 1 | 29 | Q92698 |
| High | FNSPSSPDF | 1xCarbami | 0.00245 | 0.004739 | 1 | 1 | 2 | Q92698 |
| High | FNSPSSPDF | 1xCarbami | 2.5E-06 | 0.004739 | 1 | 1 | 6 | Q92698 |
| High | CVEEDGFV | 1xCarbami | 3.49E-06 | 0.004739 | 1 | 1 | 154 | Q92698 |
| High | ELFILDEASL | 1xCarbami | 0.00092 | 0.004739 | 1 | 1 | 2 | Q92698 |
| High | ELFILDEASL | 1xCarbami | 0.000216 | 0.004739 | 1 | 1 | 11 | Q92698 |
| High | VVLVSNYTC | 1xCarbami | 0.000245 | 0.004739 | 1 | 1 | 3 | Q92698 |
| High | MSVSSLSSI | 1xCarbami | 6.57E-06 | 0.004739 | 1 | 1 | 1 | Q92698 |
| High | KPLSQLTNC | 1xCarbami | 0.116337 | 0.004739 | 1 | 1 | 1 | Q92698 |
| High | KPLSQLTNC | 1xCarbami | 5.2E-07 | 0.004739 | 1 | 1 | 27 | Q92698 |
| High | KPLSQLTNC | 1xCarbami | 1.19E-09 | 0.004739 | 1 | 1 | 119 | Q92698 |
| High | KSSSETQIQI | 1xCarbami | 0.000282 | 0.004739 | 1 | 1 | 11 | Q92698 |
| High | KSSSETQIQI | 1xCarbami | 2.31E-06 | 0.004739 | 1 | 1 | 43 | Q92698 |
| High | SSSETQIQEC | 1xCarbami | 5.36E-05 | 0.004739 | 1 | 1 | 24 | Q92698 |
| High | SSSETQIQEC | 1xCarbami | 4.94E-06 | 0.004739 | 1 | 1 | 60 | Q92698 |
| High | ALEPQLSGK |  | 0.004906 | 0.004739 | 1 | 1 | 35 | Q92698 |
| High | ALEPQLSGK |  | 0.000164 | 0.004739 | 1 | 1 | 6 | Q92698 |
| High | ALEPQLSGK |  | 0.00071 | 0.004739 | 1 | 1 | 7 | Q92698 |
| High | ALHDPLEK |  | 0.004028 | 0.004739 | 1 | 1 | 13 | Q92698 |
| High | ALHDPLEKI |  | 4.14E-08 | 0.004739 | 1 | 1 | 285 | Q92698 |
| High | ALHDPLEKI |  | 1.71E-09 | 0.004739 | 1 | 1 | 153 | Q92698 |
| High | AVVVSPPSL |  | 7.95E-05 | 0.004739 | 1 | 1 | 44 | Q92698 |
| High | AVVVSPPSL |  | 2.94E-05 | 0.004739 | 1 | 1 | 3 | Q92698 |
| High | DAAASEADI |  | 0.001249 | 0.004739 | 1 | 1 | 1 | Q92698 |
| High | DAAASEADI |  | 0.000106 | 0.004739 | 1 | 1 | 23 | Q92698 |
| High | DAAASEADI |  | 0.026255 | 0.004739 | 1 | 1 | 1 | Q92698 |
| High | DALVLYEPP |  | 5.86E-05 | 0.004739 | 1 | 1 | 68 | Q92698 |
| High | DALVLYEPP |  | 1.32E-06 | 0.004739 | 1 | 1 | 27 | Q92698 |
| High | DALVLYEPP |  | 5.06E-07 | 0.004739 | 1 | 1 | 10 | Q92698 |
| High | DEIDQKLEG |  | 6.32E-05 | 0.004739 | 1 | 1 | 11 | Q92698 |
| High | DEVLQAAW |  | 4.45E-07 | 0.004739 | 1 | 1 | 4 | Q92698 |
| High | DRPSVNSIL |  | 0.00165 | 0.004739 | 1 | 1 | 1 | Q96PY6 |
| High | EKLPVHVVA |  | 5.14E-06 | 0.004739 | 1 | 1 | 72 | Q92698 |
| High | ELFILDEASL |  | 1.5E-06 | 0.004739 | 1 | 1 | 222 | Q92698 |
| High | ELTSIVNR |  | 0.001595 | 0.004739 | 1 | 1 | 10 | Q92698 |
| High | EVAVLANM |  | 0.142523 | 0.004739 | 1 | 1 | 3 | Q96PY6 |
| High | FNSPSSPDF |  | 2.17E-06 | 0.004739 | 1 | 1 | 133 | Q92698 |
| High | GRDAAASE/ |  | 2.46E-06 | 0.004739 | 1 | 1 | 17 | Q92698 |
| High | HFELPILK |  | 3.79E-05 | 0.004739 | 1 | 1 | 176 | Q92698 |
| High | HFELPILKG |  | 0.004021 | 0.004739 | 1 | 1 | 4 | Q92698 |
| High | HFSLGELK |  | 4.79E-05 | 0.004739 | 1 | 1 | 207 | Q92698 |
| High | HFSLGELKE |  | 4.38E-08 | 0.004739 | 1 | 1 | 139 | Q92698 |
| High | HPNIVQYR |  | 0.004413 | 0.004739 | 1 | 1 | 2 | Q96PY6 |
| High | KHFELPILK |  | 0.001088 | 0.004739 | 1 | 1 | 120 | Q92698 |
| High | LDGTMSIK |  | 0.000439 | 0.004739 | 1 | 1 | 37 | Q92698 |
| High | LDGTMSIKI |  | 0.002048 | 0.004739 | 1 | 1 | 9 | Q92698 |
| High | LDGTMSIKI |  | 0.038011 | 0.004739 | 1 | 1 | 1 | Q92698 |
| High | LDKEKLPVH |  | 2.99E-08 | 0.004739 | 1 | 1 | 177 | Q92698 |
| High | LEGFMNQR |  | 0.001825 | 0.004739 | 1 | 1 | 35 | Q92698 |
| High | IGEGSFGK |  | 0.003307 | 0.004739 | 1 | 1 | 1 | Q96PY6 |
| High | LHVGVLQK |  | 0.0009 | 0.004739 | 1 | 1 | 88 | Q92698 |
| High | LKNSENQTY |  | 7.35E-06 | 0.004739 | 1 | 1 | 36 | Q92698 |
| High | LKNSENQTY |  | 0.000206 | 0.004739 | 1 | 1 | 11 | Q92698 |
| High | LLSAGTIEEK |  | 0.000128 | 0.004739 | 1 | 1 | 151 | Q92698 |
| High | LLSAGTIEEK |  | 6.68E-05 | 0.004739 | 1 | 1 | 3 | Q92698 |
| High | LPVHVVVD |  | 6.57E-06 | 0.004739 | 1 | 1 | 167 | Q92698 |
| High | LPVHVVVD |  | 0.000305 | 0.004739 | 1 | 1 | 4 | Q92698 |
| High | IQPLAIDGG |  | 0.001598 | 0.004739 | 1 | 1 | 8 | Q92698 |
| High | IQPLAIDGG |  | 8.82E-06 | 0.004739 | 1 | 1 | 89 | Q92698 |
| High | IQPLAIDGG |  | 1.11E-07 | 0.004739 | 1 | 1 | 165 | Q92698 |
| High | LRELTSIVNF |  | 0.000748 | 0.004739 | 1 | 1 | 183 | Q92698 |
| High | LTPLQTELYI |  | 0.000356 | 0.004739 | 1 | 1 | 40 | Q92698 |
| High | LTPLQTELYI |  | 2.78E-05 | 0.004739 | 1 | 1 | 94 | Q92698 |
| High | LTPLQTELYI |  | 0.219871 | 0.004739 | 1 | 1 | 1 | Q92698 |
| High | LVMFDPDV |  | 5.77E-06 | 0.004739 | 1 | 1 | 114 | Q92698 |
| High | IVQNILGNE |  | 0.024054 | 0.004739 | 1 | 1 | 2 | Q96PY6 |
| High | MLVLDYILA |  | 5.63E-06 | 0.004739 | 1 | 1 | 98 | Q92698 |
| High | MSVSSLSSI |  | 0.000209 | 0.004739 | 1 | 1 | 19 | Q92698 |
| High | MSVSSLSSI |  | 7.73E-06 | 0.004739 | 1 | 1 | 119 | Q92698 |
| High | NSENQTYQ/ |  | 7.47E-07 | 0.004739 | 1 | 1 | 47 | Q92698 |
| High | NSENQTYQ/ |  | 0.000207 | 0.004739 | 1 | 1 | 13 | Q92698 |
| High | NWYNEVGK |  | 0.005236 | 0.004739 | 1 | 1 | 88 | Q92698 |
| High | NWYNEVGK |  | 0.000543 | 0.004739 | 1 | 1 | 5 | Q92698 |
| High | QAKPAEELL |  | 5.04E-05 | 0.004739 | 1 | 1 | 119 | Q92698 |
| High | QAKPAEELL |  | 0.00054 | 0.004739 | 1 | 1 | 1 | Q92698 |
| High | QAKPAEELL |  | 0.001735 | 0.004739 | 1 | 1 | 2 | Q92698 |
| High | RALHDPLEK |  | 0.006919 | 0.004739 | 1 | 1 | 4 | Q92698 |
| High | RALHDPLEK |  | 9.54E-09 | 0.004739 | 1 | 1 | 236 | Q92698 |
| High | RSLAPSQLA |  | 9.69E-05 | 0.004739 | 1 | 1 | 10 | Q92698 |
| High | RSLAPSQLA |  | 0.121563 | 0.004739 | 1 | 1 | 1 | Q92698 |
| High | RVLISGTPIC |  | 0.001506 | 0.004739 | 1 | 1 | 32 | Q92698 |
| High | RYLYVR |  | 0.005017 | 0.004739 | 1 | 1 | 82 | Q92698 |
| High | SLAPSQLAK |  | 0.002679 | 0.004739 | 1 | 1 | 38 | Q92698 |
| High | SLAPSQLAK |  | 0.013897 | 0.004739 | 1 | 1 | 3 | Q92698 |
| High | SILSKPFK |  | 0.001511 | 0.004739 | 1 | 1 | 30 | Q92698 |
| High | SILSKPFKVF |  | 1.49E-07 | 0.004739 | 1 | 1 | 129 | Q92698 |
| High | SSDKVVLVS |  | 1.28E-06 | 0.004739 | 1 | 1 | 265 | Q92698 |
| High | TSDILSK |  | 0.000624 | 0.004739 | 1 | 1 | 11 | Q92698 |
| High | TSDILSKYLP |  | 0.001429 | 0.004739 | 1 | 1 | 4 | Q92698 |
| High | VLISGTPIQ |  | 9.12E-09 | 0.004739 | 1 | 1 | 38 | Q92698 |
| High | VLISGTPIQ |  | 1.38E-08 | 0.004739 | 1 | 1 | 70 | Q92698 |
| High | VPIPNYQGF |  | 4.87E-05 | 0.004739 | 1 | 1 | 70 | Q92698 |
| High | VSSPILIISE |  | 4.33E-06 | 0.004739 | 1 | 1 | 209 | Q92698 |
| High | VSSPILIISE |  | 0.023489 | 0.004739 | 1 | 1 | 46 | Q92698 |
| High | VVERFNPS |  | 0.028862 | 0.004739 | 1 | 1 | 1 | Q92698 |
| High | VVLVSNYTC |  | 4.09E-06 | 0.004739 | 1 | 1 | 179 | Q92698 |
| High | WGLRDEVL |  | 0.009857 | 0.004739 | 1 | 1 | 1 | Q92698 |

| # Missed Cl | Theo. MH+ | Confidence | Percolator | Percolator | XCorr (by Search Engine): Sequest HT |
| --- | --- | --- | --- | --- | --- |
| 0 | 2251.975 | High | 0.000146 | 1.1E-12 | 2.59 |
| 0 | 1063.502 | High | 0.000146 | 3.86E-10 | 2.22 |
| 1 | 2960.504 | High | 0.000146 | 0.000185 | 0.78 |
| 2 | 3868.856 | High | 0.000146 | 6.31E-16 | 4.02 |
| 1 | 3740.761 | High | 0.000146 | 6.31E-16 | 4.44 |
| 0 | 2807.155 | High | 0.000146 | 7.31E-15 | 6.22 |
| 0 | 2727.189 | High | 0.000146 | 6.31E-16 | 7.75 |
| 1 | 1663.866 | High | 0.000146 | 5.17E-11 | 2.79 |
| 0 | 1486.774 | High | 0.000146 | 0.000872 | 0.64 |
| 0 | 1089.44 | High | 0.000146 | 5.74E-06 | 1.33 |
| 0 | 1140.555 | High | 0.000146 | 7.07E-10 | 2.32 |
| 1 | 1280.672 | High | 0.000146 | 0.000322 | 1.97 |
| 0 | 1940.864 | High | 0.000146 | 1.7E-07 | 2.53 |
| 0 | 994.4969 | High | 0.000146 | 2.78E-06 | 1.89 |
| 1 | 1150.598 | High | 0.000146 | 2.83E-07 | 3.12 |
| 0 | 999.5275 | High | 0.000146 | 5.15E-09 | 1.93 |
| 0 | 1477.72 | High | 0.000146 | 5.61E-06 | 2.71 |
| 1 | 2378.263 | High | 0.000146 | 6.78E-11 | 4.92 |
| 0 | 944.4159 | High | 0.000146 | 6.5E-07 | 2.09 |
| 1 | 2262.04 | High | 0.000146 | 5.83E-08 | 3.36 |
| 2 | 2418.141 | High | 0.000146 | 8.77E-07 | 4.91 |
| 1 | 1547.775 | High | 0.000146 | 3.53E-08 | 2.31 |
| 0 | 2020.861 | High | 0.000146 | 1.45E-11 | 4.73 |
| 1 | 2176.962 | High | 0.000146 | 3.37E-11 | 5.02 |
| 0 | 2235.98 | High | 0.000146 | 2.2E-13 | 4.2 |
| 0 | 880.4445 | High | 0.000146 | 7.02E-09 | 1.84 |
| 1 | 1008.539 | High | 0.000146 | 3.62E-09 | 2.37 |
| 0 | 1010.472 | High | 0.000146 | 5.83E-08 | 2.12 |
| 2 | 2818.404 | High | 0.000146 | 7.81E-16 | 4.53 |
| 0 | 1422.803 | High | 0.000146 | 4.82E-15 | 2.59 |
| 0 | 1355.709 | High | 0.000146 | 1.9E-12 | 3.38 |
| 1 | 1483.804 | High | 0.000146 | 1.84E-15 | 4.28 |
| 2 | 3065.655 | High | 0.000146 | 0.00013 | 1.63 |
| 0 | 1805.841 | High | 0.000146 | 1.06E-11 | 3.26 |
| 1 | 1738.806 | High | 0.000146 | 4.96E-08 | 3.2 |
| 1 | 2346.31 | High | 0.000146 | 1.24E-05 | 0.87 |
| 1 | 1610.8 | High | 0.000146 | 9.81E-10 | 2.93 |
| 1 | 1960.904 | High | 0.000146 | 6.31E-16 | 5.95 |
| 1 | 1944.909 | High | 0.000146 | 6.31E-16 | 5.9 |
| 0 | 1398.679 | High | 0.000146 | 4.5E-11 | 3.78 |
| 1 | 1639.858 | High | 0.000146 | 1.83E-10 | 2.43 |
| 1 | 1928.914 | High | 0.000146 | 6.31E-16 | 5.66 |
| 0 | 1804.803 | High | 0.000146 | 2.05E-09 | 2.87 |
| 0 | 1788.808 | High | 0.000146 | 1.52E-14 | 3.88 |
| 0 | 1772.813 | High | 0.000146 | 6.31E-16 | 5.49 |
| 1 | 1863.875 | High | 0.000146 | 1.23E-13 | 5.14 |
| 0 | 1815.747 | High | 0.000146 | 1.72E-08 | 3.29 |
| 0 | 1735.78 | High | 0.000146 | 1.88E-14 | 3.73 |
| 1 | 2647.267 | High | 0.000146 | 1.15E-11 | 3.56 |
| 0 | 1197.572 | High | 0.000146 | 4.24E-12 | 1.96 |
| 1 | 1353.673 | High | 0.000146 | 1.09E-05 | 1.5 |
| 0 | 1330.631 | High | 0.000146 | 9.08E-12 | 2.86 |
| 1 | 2397.269 | High | 0.000146 | 3.04E-12 | 2.78 |
| 0 | 1664.886 | High | 0.000146 | 6.31E-12 | 2.56 |
| 0 | 1272.648 | High | 0.000146 | 2.52E-13 | 3.46 |
| 1 | 3473.615 | High | 0.000146 | 4.5E-07 | 1.76 |
| 0 | 1648.891 | High | 0.000146 | 4.04E-15 | 2.95 |
| 1 | 1471.773 | High | 0.000146 | 2.88E-14 | 4.49 |
| 1 | 1003.503 | High | 0.000146 | 1.08E-09 | 2.42 |
| 0 | 1854.736 | High | 0.000146 | 2.13E-06 | 3.13 |
| 0 | 1343.678 | High | 0.000146 | 1.61E-12 | 3.91 |
| 0 | 1774.77 | High | 0.000146 | 1.73E-15 | 4.38 |
| 0 | 875.408 | High | 0.000146 | 3.14E-08 | 1.09 |
| 1 | 3059.471 | High | 0.000146 | 2.02E-08 | 1.18 |
| 1 | 3043.476 | High | 0.000146 | 1.55E-15 | 3.72 |
| 0 | 2416.101 | High | 0.000146 | 3.4E-15 | 4.99 |
| 1 | 2507.138 | High | 0.000146 | 1.96E-09 | 4.92 |
| 1 | 2427.172 | High | 0.000146 | 6.2E-11 | 3.93 |
| 1 | 2298.216 | High | 0.000146 | 8.44E-11 | 3.95 |
| 2 | 2792.469 | High | 0.000146 | 1.53E-14 | 5.08 |
| 0 | 2826.268 | High | 0.000146 | 0.00025 | 1.22 |
| 0 | 2746.302 | High | 0.000146 | 6.31E-16 | 5.13 |
| 0 | 2666.336 | High | 0.000146 | 6.31E-16 | 7.06 |
| 1 | 2123.947 | High | 0.000146 | 1.17E-10 | 4.61 |
| 1 | 2043.98 | High | 0.000146 | 1.27E-15 | 5.02 |
| 0 | 1995.852 | High | 0.000146 | 2.26E-12 | 4.04 |
| 0 | 1915.885 | High | 0.000146 | 7.81E-15 | 3.28 |
| 0 | 942.5255 | High | 0.000146 | 1.06E-07 | 1.93 |
| 1 | 2330.315 | High | 0.000146 | 3.26E-11 | 2.8 |
| 2 | 2573.448 | High | 0.000146 | 1.05E-09 | 2.82 |
| 0 | 922.4993 | High | 0.000146 | 6.6E-08 | 2.47 |
| 1 | 2909.541 | High | 0.000146 | 6.31E-16 | 7.56 |
| 2 | 3265.747 | High | 0.000146 | 6.31E-16 | 7 |
| 0 | 1085.656 | High | 0.000146 | 5.78E-12 | 2.36 |
| 1 | 2076.112 | High | 0.000146 | 5.45E-13 | 4.28 |
| 0 | 905.3959 | High | 0.000146 | 4.05E-09 | 1.99 |
| 1 | 1617.746 | High | 0.000146 | 1.16E-11 | 3.81 |
| 2 | 1886.931 | High | 0.000146 | 6E-06 | 2.18 |
| 0 | 2006.059 | High | 0.000146 | 2.8E-12 | 3.17 |
| 1 | 2362.265 | High | 0.000146 | 6.31E-16 | 3.91 |
| 2 | 2619.403 | High | 0.000146 | 6.31E-16 | 5.69 |
| 1 | 1722.812 | High | 0.000146 | 3.35E-12 | 1.94 |
| 0 | 2476.226 | High | 0.000146 | 6.31E-16 | 5.07 |
| 0 | 1257.68 | High | 0.000146 | 7.87E-09 | 2.52 |
| 1 | 1673 | High | 0.000146 | 8.58E-15 | 3.32 |
| 0 | 1860.897 | High | 0.000146 | 6.31E-16 | 4.19 |
| 0 | 931.5207 | High | 0.000146 | 7.25E-09 | 1.99 |
| 0 | 974.5339 | High | 0.000146 | 0.000425 | 1.23 |
| 0 | 1789.847 | High | 0.000146 | 1.1E-15 | 2.37 |
| 2 | 1830.869 | High | 0.000146 | 1.49E-15 | 3.79 |
| 0 | 996.5877 | High | 0.000146 | 9.92E-13 | 2.6 |
| 1 | 1209.71 | High | 0.000146 | 6.58E-08 | 2.99 |
| 0 | 930.5043 | High | 0.000146 | 1.74E-12 | 2.6 |
| 1 | 2772.384 | High | 0.000146 | 6.31E-16 | 5.33 |
| 0 | 1026.548 | High | 0.000146 | 8.22E-08 | 1.74 |
| 1 | 1124.683 | High | 0.000146 | 2.92E-09 | 2.18 |
| 0 | 864.4495 | High | 0.000146 | 3.37E-10 | 2.05 |
| 1 | 992.5445 | High | 0.000146 | 1.32E-08 | 1.66 |
| 2 | 1148.646 | High | 0.000146 | 1.49E-05 | 1.31 |
| 2 | 2029.206 | High | 0.000146 | 6.31E-16 | 7.2 |
| 0 | 994.4775 | High | 0.000146 | 9.99E-09 | 2.04 |
| 0 | 794.4043 | High | 0.000146 | 4.12E-08 | 1.3 |
| 0 | 893.5567 | High | 0.000146 | 1.86E-09 | 2.08 |
| 1 | 2182.073 | High | 0.000146 | 2.01E-14 | 5.76 |
| 2 | 2338.175 | High | 0.000146 | 5.57E-11 | 4.39 |
| 0 | 1060.588 | High | 0.000146 | 1.79E-11 | 1.45 |
| 1 | 1604.901 | High | 0.000146 | 3.82E-12 | 3.8 |
| 0 | 1415.862 | High | 0.000146 | 1.54E-14 | 4.2 |
| 1 | 2302.387 | High | 0.000146 | 1.41E-10 | 2.19 |
| 0 | 1098.615 | High | 0.000146 | 7.29E-09 | 2.17 |
| 1 | 1826.949 | High | 0.000146 | 3.09E-14 | 3.5 |
| 2 | 2802.409 | High | 0.000146 | 6.31E-16 | 5.64 |
| 1 | 1200.706 | High | 0.000146 | 1.19E-09 | 3.32 |
| 0 | 1205.678 | High | 0.000146 | 2.05E-10 | 1.75 |
| 1 | 1361.779 | High | 0.000146 | 4.74E-13 | 3.24 |
| 2 | 1778.032 | High | 0.000146 | 0.001359 | 1.3 |
| 0 | 2219.985 | High | 0.000146 | 1.13E-14 | 3.4 |
| 0 | 1877.003 | High | 0.000146 | 4.86E-06 | 1.35 |
| 0 | 1406.808 | High | 0.000146 | 1.07E-14 | 2.84 |
| 0 | 1339.714 | High | 0.000146 | 5.74E-11 | 2.95 |
| 1 | 1467.809 | High | 0.000146 | 2.26E-14 | 4.31 |
| 0 | 1940.894 | High | 0.000146 | 6.31E-16 | 3.55 |
| 1 | 2096.996 | High | 0.000146 | 5.61E-11 | 4.6 |
| 0 | 1009.474 | High | 0.000146 | 1.24E-07 | 1.75 |
| 1 | 1578.781 | High | 0.000146 | 5.6E-10 | 2.26 |
| 0 | 1312.711 | High | 0.000146 | 1.96E-12 | 3.42 |
| 1 | 2633.407 | High | 0.000146 | 5.51E-10 | 3.62 |
| 2 | 2761.502 | High | 0.000146 | 8.85E-09 | 3.66 |
| 1 | 1078.6 | High | 0.000146 | 2.41E-07 | 2.31 |
| 2 | 3065.642 | High | 0.000146 | 6.31E-16 | 5.9 |
| 1 | 1070.632 | High | 0.000146 | 9.28E-12 | 2.74 |
| 2 | 1226.733 | High | 0.000146 | 0.000281 | 1.13 |
| 1 | 3915.08 | High | 0.000146 | 6.32E-09 | 1.88 |
| 1 | 869.4992 | High | 0.000146 | 1.12E-07 | 1.39 |
| 0 | 914.5306 | High | 0.000146 | 2.5E-08 | 1.7 |
| 2 | 1637.945 | High | 0.000146 | 1.28E-06 | 2.07 |
| 0 | 919.5611 | High | 0.000146 | 6.38E-09 | 2.21 |
| 1 | 2298.297 | High | 0.000146 | 6.31E-16 | 4.59 |
| 1 | 2286.186 | High | 0.000146 | 6.31E-16 | 3.75 |
| 0 | 763.4196 | High | 0.000146 | 7.78E-10 | 1.89 |
| 1 | 1363.783 | High | 0.000146 | 5.6E-09 | 3.34 |
| 0 | 3758.979 | High | 0.000146 | 6.31E-16 | 7.77 |
| 1 | 3887.074 | High | 0.000146 | 6.31E-16 | 7.48 |
| 0 | 1397.754 | High | 0.000146 | 1.8E-12 | 2.17 |
| 0 | 1624.894 | High | 0.000146 | 5.7E-15 | 2.46 |
| 1 | 2499.433 | High | 0.000146 | 4.57E-06 | 0.98 |
| 1 | 2273.127 | High | 0.000146 | 7.56E-06 | 3.16 |
| 0 | 1869 | High | 0.000146 | 4.99E-15 | 4.39 |
| 1 | 2988.512 | High | 0.000146 | 5.62E-07 | 1.46 |

